# Multimodal Mitigations for Cybersickness in Motion Base Simulators

**DOI:** 10.1101/2023.12.11.570846

**Authors:** Séamas Weech, Anouk Lamontagne

## Abstract

1.

**Background:** Virtual reality (VR) technologies that integrate with motion-base simulators (MBS) have the potential to accelerate personnel training and enhance workplace safety. Motion sickness on an MBS is a widespread problem with vast individual differences that are likely related to idiosyncrasies in estimates of head, body, and vehicle motions. When combined with head-mounted VR, we term the emergent symptoms ‘cybersickness’.

**Methods:** We conducted two experiments that evaluated cybersickness mitigations in an MBS. In Experiment 1 (N = 8), we tested the effectiveness of a light-touch body harness attached to a mobile-elevated work platform (MEWP) simulator during two nauseogenic VR tasks. In Experiment 2 (N = 14, 7 of whom completed Experiment 1), we tested the effectiveness of a dynamic field-of-view (dFOV) modifier that adaptively restricted the FOV for vehicle rotations in the same VR tasks. We gathered subjective sickness data and qualitative evaluations of the mitigations after the fact.

**Results:** We observed a reduced level of sickness in both Experiment 1 and 2 when mitigations were applied. In Experiment 1, the use of a harness led to a mild decrease in total cybersickness of between 3-11%, which was only significant for the nausea dimension. In Experiment 2, the use of dFOV imparted a large benefit to comfort, up to a 45% improvement. Both mitigations primarily improved comfort in a bumpy trench traversal task.

**Conclusions:** Cybersickness mitigations can help to deliver VR training for longer, and to more users. The type of content undertaken should be considered when employing new mitigations.

## 2. Introduction

Veridical self-motion perception requires humans to constantly update an estimate of the position and orientation of the head and body in space. This task must be performed while integrating across several distinct reference frames that are necessary for a variety of motor control tasks to be executed (e.g., head-centered, eye-centered, and body-centered reference frames, Murdison et al. 2017). Given the multitude of noisy and high-bandwidth information arriving at integration sites of the brain every moment, the level of challenge posed by this task is considerable. Regardless, in everyday life we perform well, with this system entering very few failure states. This is possible given that the key neural processes subserving self-motion have been formed through continual exposure to a natural world that conforms to stable physics and immutable rules, meaning that it operates well within the confines of these rules.

When virtual reality (VR) replaces the stable, immutable physics of real reality, this system is flung into disarray. Many actions are left intact, but at the level of unconscious processes like the modality cue-conflict resolution system in the brainstem, the transition from real reality to VR generates negative effects. Most pronounced of these is cybersickness. Cybersickness impacts the well-being, physical coordination, and visual processing of VR users (Stanney et al. 2020). This problem severely limits the full capabilities of the technology and creates the need for powerful yet lightweight mitigations. An example of settings where cybersickness mitigations are sought after is the domain of operational training, where novice personnel can replace hours of practice on expensive and potentially dangerous machinery with high-fidelity simulator training hours. In these settings, using a motion base to deliver inertial motion cues that are consistent with vehicle physics offers an effective method to reduce cybersickness (Blana 1996; Klüver et al. 2015). For simulator-based skills acquisition, cybersickness must be avoided as best as possible to maximize the duration and frequency of operational training sessions.

For cybersickness mitigations to be implemented effectively in the field, there must be two aspects that are welldefined: (a) The impact of the mitigation on user comfort, and (b) the associated costs of implementing the mitigation. In this research, we set out to define the former (a) part of the equation.

We elected to examine two mitigations that target self-motion perception. Uncertainty in the self-motion and orientation resolution process is a key contributor to cybersickness (Bos et al. 2008; Weech et al. 2019). Specifically, it is thought that cybersickness relates to the multisensory re-weighting process that is dedicated to resolving this uncertainty and providing a stable state estimate which subserves smooth control of behavior in complex environments (Weech et al. 2018b, 2020a). This theory gains support from reports that cybersickness is significantly reduced when multisensory stimulation is used to bias the process of self-motion and orientation estimation (Bos 2015; Weech et al. 2018a, 2020c).

First, we test the use of a harness attached to the frame of the motion base simulator (MBS). This harness provides minimal support for balance yet offers subtle haptic cues to the user about where their body is located in 3D space—the estimation of which is not trivial when performing in the non-zero-latency world of VR (Palmisano et al. 2020). Based on evidence that light haptic stimulation forms a valuable source of information for feedback-control of body posture (Jeka 1997; Krishnamoorthy et al. 2002; Lackner et al. 2001; Rogers et al. 2001), we postulated that a body harness tethered loosely to the physical frame of the simulator would reduce uncertainty in self-motion perception and ameliorate discomfort as a result.

Second, we assess a dynamic field-of-view (dFOV) limiter that contrast-modulates the eccentric field-of-view when the user’s vehicle rotates in virtual space. Similar techniques have been used before in experimental (Fernandes & Feiner 2016) and commercial settings (e.g., Google Earth VR). Cue-conflict signals that accumulate in the brain’s emesis centers (a brainstem-cerebellum loop, Yates et al. 2014) primarily arise from rotation epochs (Oman & Cullen 2014; Weech et al. 2018a). Models of cybersickness reveal that optic flow directly correlates with nausea (Lee et al. 2017), and dFOV limits flow during these sensitive epochs of imposed self-motion. Given the high prominence of peripheral optic flow in the self-motion system (Gibson 1961; Warren & Hannon 1988), we postulated that a dFOV modifier would limit cue-conflict accumulation and reduce cybersickness.

Here, we designed a study to explore and verify the utility of two cybersickness mitigations during VR tasks on an MBS: A harness attached to the frame of the simulator, and a visual dFOV modifier. We captured sickness scores through self-reported subjective measures using a gold-standard questionnaire. Where possible, we compared sickness where a mitigation was used versus no mitigation (control conditions) and examined whether mitigations specifically affected sub-types of cybersickness (nausea, disorientation, and oculomotor discomfort).

## 3. Methods

### 3.1. Participants

We recruited a convenience sample of healthy adults from the workplace population of a local technology company and acquaintances. All participants were asked if they owned a VR HMD (none did). Sample size was determined by practicality. In Experiment 1, N = 8; for Experiment 2, N = 14. Seven participants completed both experiments. All individuals were reliably informed that participation is voluntary and that they were not obliged to take part in the research if they did not wish to do so. Participants were not compensated for their time. The study ethics protocol was approved by the McGill University Research Ethics Board and was conducted according to ethical principles stated in the Declaration of Helsinki (2013).

### 3.2. Procedure

The letter of information, consent, and obtention of demographic information took place upon the participant’s arrival to the study location. Upon arrival, participants were reminded about the general goal of the research, the requirements of the VR task, and the risks of the experiment.

Following this, participants were informed about the task to be conducted in virtual reality. The task was conducted within a virtual environment that depicted a 3D image of an industrial workplace site (Fig 1), rendered on a head mounted display (Oculus Rift S, Facebook, Menlo Park, CA, Fig 2) that was driven by a high-end notebook computer (Lenovo P71, 24GB RAM, i7-7700HQ processor, Quadro P4000 GPU). The participant was positioned on a motion base simulator (Mobile Elevated Work Platform simulator, MEWP, Serious Labs Inc., Edmonton, AB, Canada; Fig 2) that mimicked the motions and vibrations of a MEWP. The participant was instructed to use handheld levers on the MEWP to drive the vehicle around a simulated environment to specified locations.

**Fig 1.**
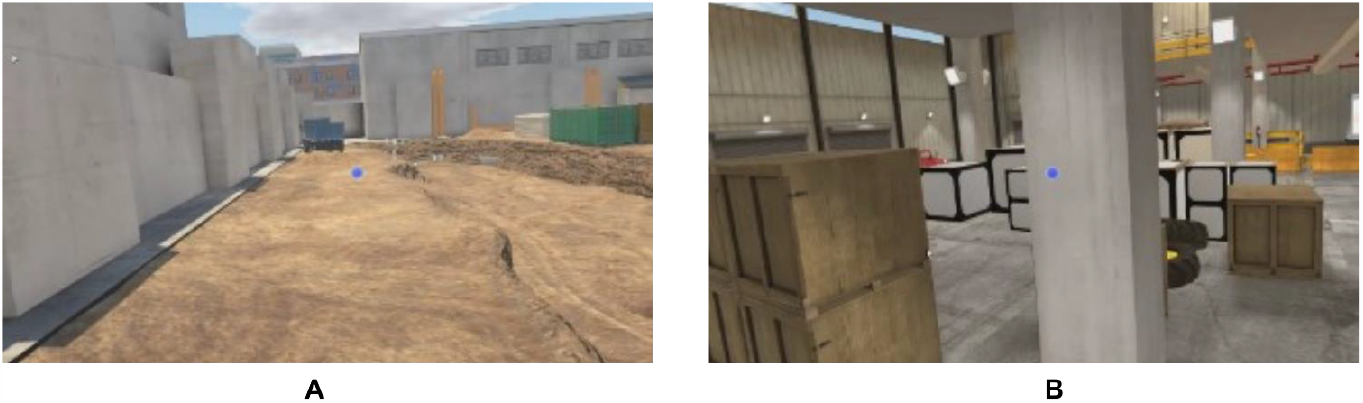
The Trench Traversal (A) and Choose Incline (B) scenarios used as tasks in the current studies.

**Fig 2.**
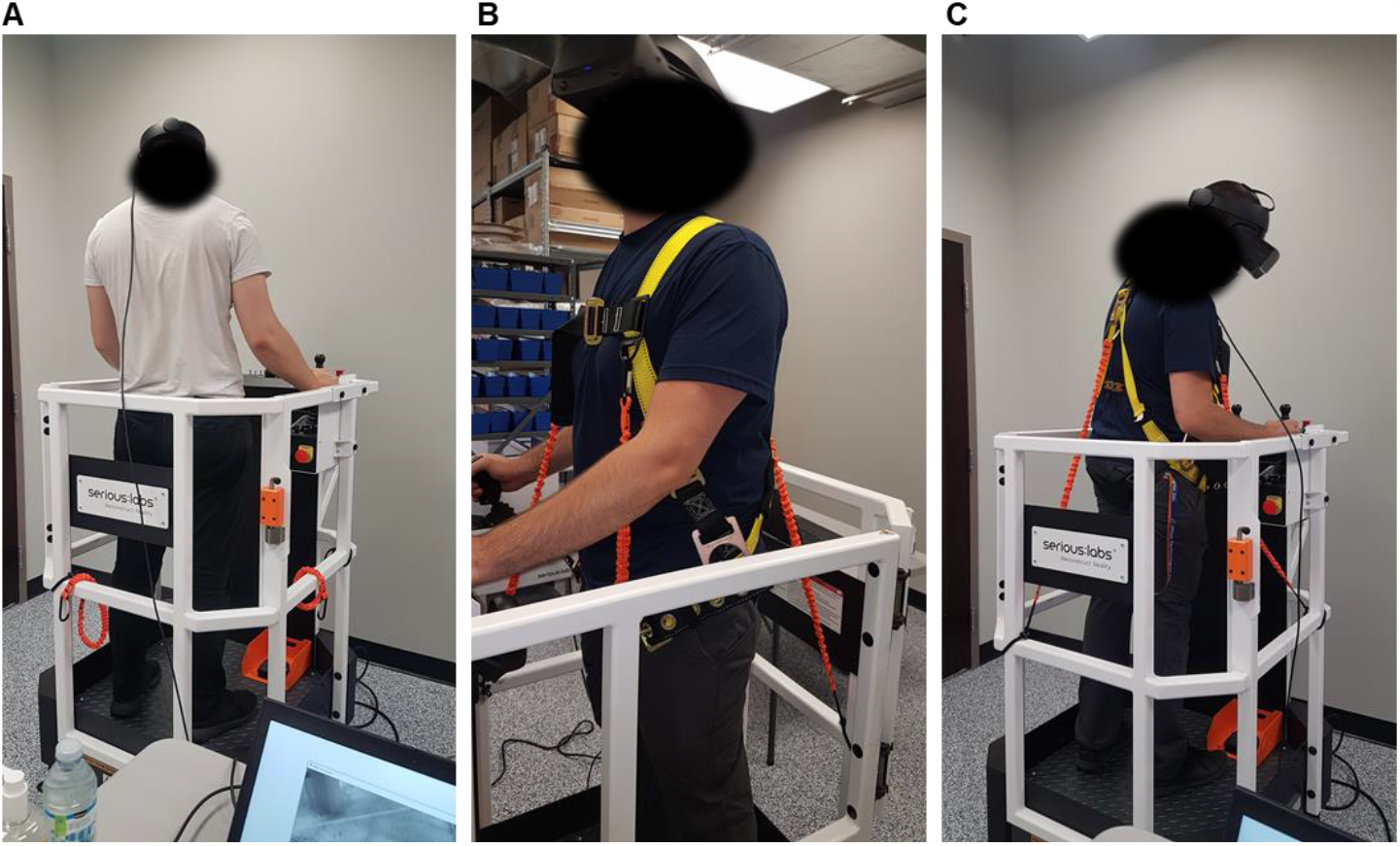
(A) The MEWP simulator in use by a participant in standard conditions. (B) A participant using the Harness in Experiment 1. (C) Reverse view of the Harness in use.

Participants were informed that the motion base will move in tandem with the virtual vehicle while the task was being completed. For each task, participants were given on-screen instructions that depended on the specific task (e.g., “take the vehicle to the glowing zone”). They were asked to perform the task as quickly as possible but without making collisions. A given task continued until the task was completed (around 5-10 min total on average). Upon a major error (e.g., a collision) the task re-started and continued until it was completed successfully. Two scenarios were undertaken: the ‘Rough Terrain Scissor Lift: Trench Traversal’ task, and the ‘Slab Scissor Lift: Choose Incline’ task (hereafter known as Trench Traversal, or TT, and Choose Incline, or CI). Note that experimental rigor necessitated an impoverished onboarding experience for participants when compared to the usual onboarding of a simulator user. As such, many comfort-related factors were intentionally neglected here (e.g., context setting, anxiety reduction) that would have resulted in elevated levels of cybersickness when compared to typical use cases with simulators such as these (for a discussion of the relevance of such factors, see Weech et al. 2019).

After each task, participants were asked to report their sickness level with the 16-item Simulator Sickness Questionnaire (SSQ; Kennedy et al. 1993). Participants took a short break in-between trials (∼2 min on average, as needed for participants to indicate that they had regained a baseline level of comfort).

In Experiment 1, participants conducted two sessions on separate days; in one session they were required to wear a harness attached to the MEWP at 3 places with elasticated lanyard tethers (Fig 2B-C, two attached to O-rings at the sides around the pectorals, and one at the upper back); the other session was a control (no harness). The order of the two sessions was counterbalanced to prevent order effects. The delivery of CI and TT tasks was also counterbalanced such that CI was the first task completed in one session and TT was the first to be completed in the other session. The harness was not intended to support stability (i.e., it was not tightly tethered), but to provide continuous self-motion information through haptic cues. In Experiment 2, participants were instructed that a dynamic field-of-view (dFOV) limiter would restrict their peripheral vision during rotations of the vehicle (Fig 3).

**Fig 3.**
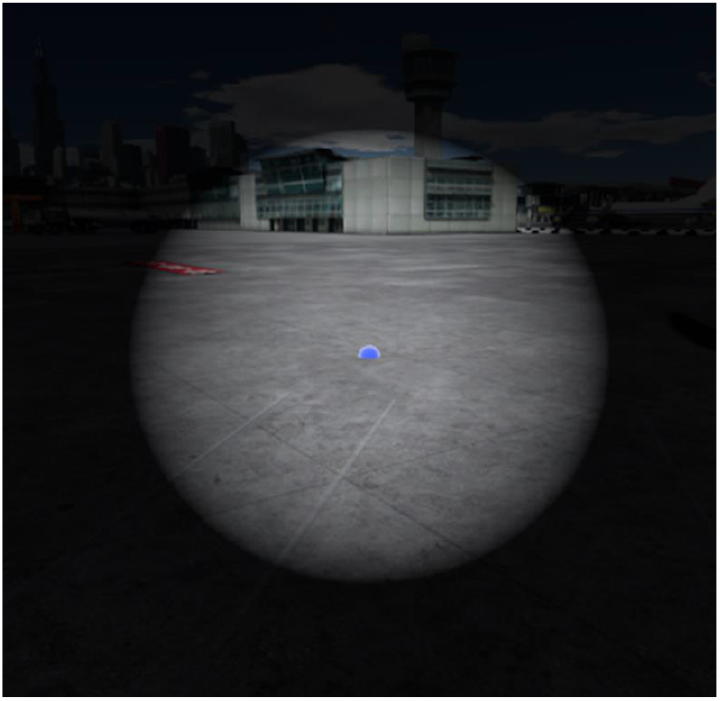
Depiction of the dynamic field-of-view (dFOV) limiter employed in Experiment 2. In the HMD, the dFOV limiter appeared less intense in contrast than in the screenshot shown here.

## 4. Results and Discussion

### 4.1. Experiment 1: Harness

The effect of wearing a harness on sickness scores is shown in Figure 4. For total SSQ scores, we documented a reduction in average sickness of 11% for the most posture-disturbing task (Trench Traversal, TT) and 3% for the other task (Choose Incline, CI). For the former, wearing a harness brought participants under the threshold for being ‘well’ on average (as per Stanney et al. 1997). These differences did not emerge as significant at the .05 alpha level (CI: *p* = .811, TT: *p* = .559), although the practical constraints that necessitated a small sample size led to a limited statistical power.

**Fig 4.**
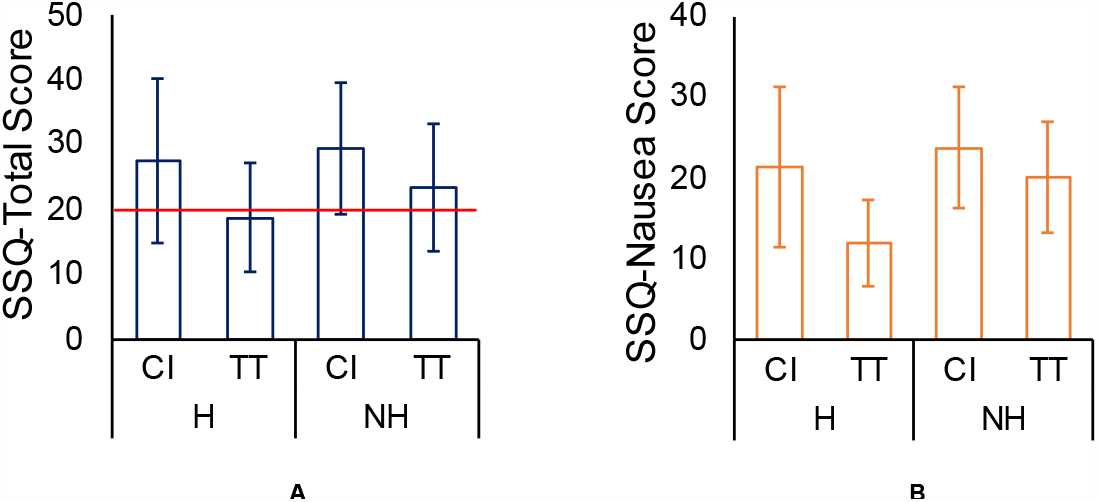
(A) Mean Simulator Sickness Questionnaire (SSQ) total scores for the two scenarios, Choose Incline (CI) and Trench Traversal (TT), across Harness (H) and No Harness (NH) conditions. Red line in (A) indicates a common threshold of ‘sick’ versus ‘well’ for total SSQ scores (Stanney et al. 1997). (B) Same plot for SSQ-Nausea, where the effects of a Harness were largest among the three SSQ subscale scores. Error bars are standard errors.

When we analyze the SSQ subscales, we see a very limited effect of harness-wearing on Disorientation (-4% detriment to sickness for CI, 4% benefit for TT), a slight difference for Oculomotor sickness (2% and 10% improvement in sickness for CI and TT respectively), and a more powerful effect on Nausea (5% improvement for CI, and a 26% improvement for TT; see Fig 4 and 5).

**Fig 5.**
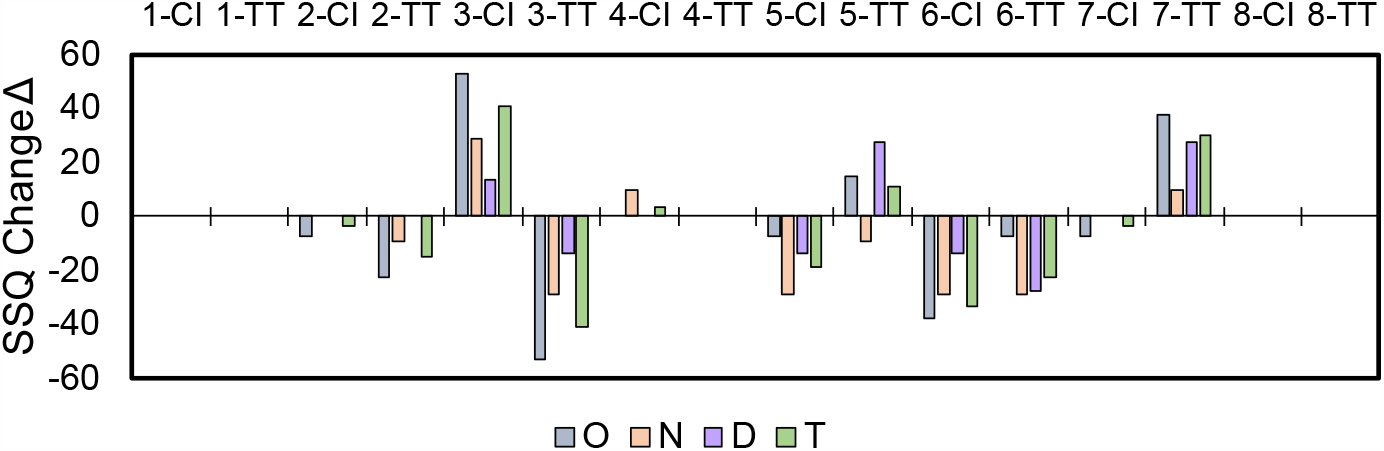
Change in SSQ total scores for the Harness task compared to control. Negative scores mean less sickness for Harness than control. CI = Choose Incline; TT = Trench Traversal. O N D T = Oculomotor, Nausea, Disorientation, Total subscale scores on the SSQ.

When assessing statistical differences among the subscale scores, the only significant difference that emerged was for the SSQ-Nausea subscale (*t*(7) = 2.55, *p* = .038, Cohen’s *d* = 0.90). This result should be considered in the context of the fact that it was not a planned comparison; we had no a-priori reason to believe that nausea would be specifically affected by the harness.

Qualitative responses (some translated from French) indicated that most participants (10) preferred having the harness on. The others indicated that they did not prefer the harness for different reasons: either “I kept being pulled by it”; “It felt too tight”; the other two did not state a reason for preferring no harness. Those that did like the harness indicated that they felt “grounded” or that they felt more “safe, like I was not going to fall off” the virtual vehicle. Others did not provide reasons for preferring the harness.

#### Experiment 1 Discussion

We found that wearing a harness in a “bumpy” VR task—Trench Traversal—brought participants under the threshold of 20 for being ‘well’ on average (Stanney et al. 1997; Weech et al., 2018a). A less “bumpy” task did not benefit much, if at all, from harness-wearing. Scores mostly improved on the nausea scale, whereas other subscales (including disorientation) did not change. This partly supports our hypothesis that the harness contributed to uncertainty resolution in self-motion estimation, which has been ascribed as a cybersickness trigger; however, it is notable that there was no significant difference between harness and control conditions for ‘total sickness scores’, which we identified a-priori as our primary statistical comparison.

We expected that the benefit of a harness would take effect at the level of unconscious body stabilization processes (Jeka 1997; Krishnamoorthy et al. 2002; Lackner et al. 2001; Rogers et al. 2001). While we made no a-priori predictions about which subscale might benefit the most, it seems reasonable to expect that disorientation would have been ameliorated by this mitigation. The fact that nausea was mostly improved might be interpreted as evidence that the benefit emerges at the level of motion integration in emetic centers of the brain (vestibular brainstem; caudal medulla, Yates et al. 2014), which transmit corollaries of the emesis response downstream to perceptual centers of the brain. No effect of a harness was seen on disorientation, which suggests that if self-motion perception was aided through the application of the harness, this aid was not consciously detected by participants.

### 4.2. Experiment 2: Dynamic Field-of-View

As for the harness mitigation, the data indicated that dFOV reduced average sickness to a level that lay below a common threshold for feeling ‘well’ (Fig 6). This was specifically the case for participants who happened to perform both the dFOV and the control task (i.e., no dFOV). However, for those who performed only the dFOV task, scores were almost precisely on the threshold—that is, they could not be classified as ‘sick’ or ‘well’ on average, according to the standard threshold of 20 on the SSQ-T (Stanney et al. 1997). As stated above, these were participants who could not complete the control task due to practical constraints related to scheduling.

**Fig 6.**
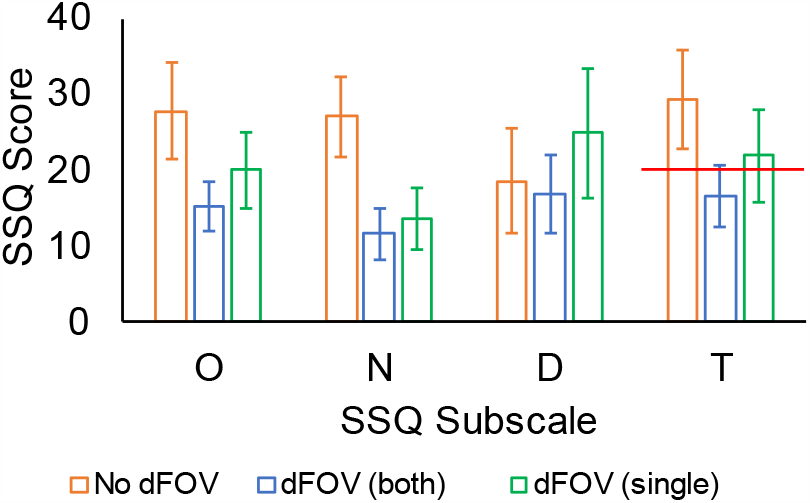
Mean SSQ scores on each subscale for the dynamic field-of-view (dFOV) task compared to control. Red line indicates a common threshold for being ‘well’ vs. ‘sick’, as in Stanney et al. (1997). Orange = No dFOV (control). Blue = dFOV (score on dFOV for participants who did both dFOV and control tasks, depicted in Fig 6 below). Green = dFOV (participants who did only dFOV tasks, not shown in Fig 6). Error bars are standard errors.

Inspecting the change in sickness from no-dFOV to dFOV for the 7 participants who completed both dFOV and control tasks, we can see a clear trend for reduced sickness when the dFOV mitigation was implemented (Fig 7). We observed a decrease in sickness of 32% (Incline) or 58% (Trench), which reduced to 17% and 45% respectively when one potential outlier participant was removed (P2 in Fig 7; Tukey’s (1977) 1.5 IQR method).

**Fig 7.**
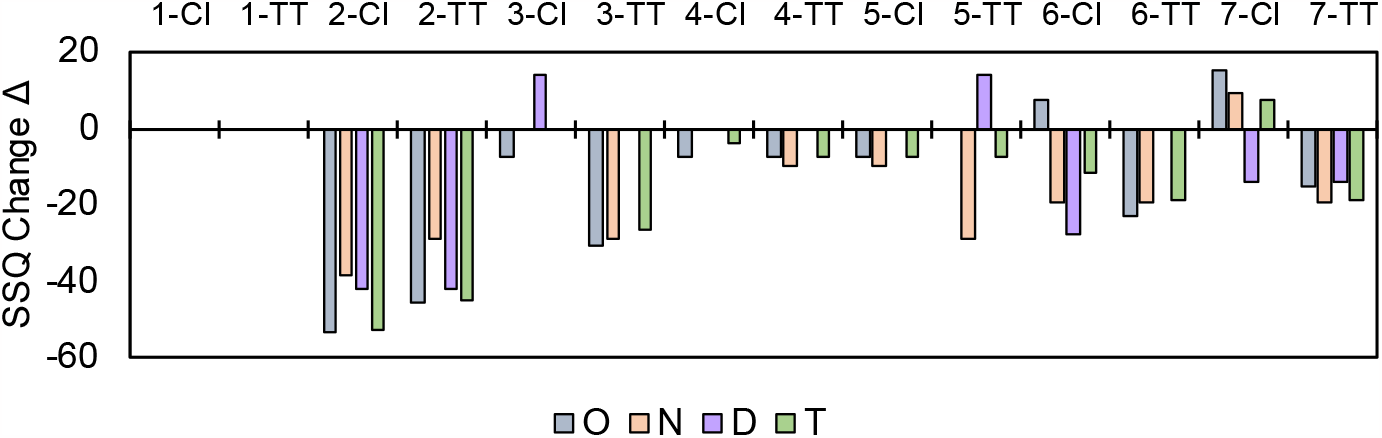
Change in SSQ total scores for the dynamic field-of-view (dFOV) task compared to control, for Participants 1-7. Negative scores mean less sickness for dFOV than control. CI = Choose Incline; TT = Trench Traversal. O N D T = Oculomotor, Nausea, Disorientation, Total subscale scores on the SSQ.

A paired-samples *t*-test (excluding the outlier) revealed a significant difference in sickness for the TT condition (*t*(5) = 3.31, *p* = .021, Cohen’s *d* = 1.35), but not for the CI condition (*t*(5) = 0.93, *p* = .39, Cohen’s *d* = 0.38)

Qualitative responses (some translated from French) indicated that the dFOV limiter was sometimes perceived as “distracting when trying to drive”; one indicated it made them “feel claustrophobic like the room was becoming smaller”. Two indicated that they “liked it”; one “did not notice it”; and the rest did not remark on it or saw no qualitative effects of dFOV.

#### Experiment 2 Discussion

We found that using a dFOV to limit the peripheral visual field improved comfort drastically in some cases, resulting in a group of participants that fell beneath the threshold for feeling ‘well’. Although participants may have felt slightly more disorientated or distracted, the level of nausea when using dFOV was much improved. The findings support the idea that dFOV limits optical flow cues that contribute to the accumulation of sensory conflict cues which drive cybersickness.

The increase in comfort was greatest once again for the “bumpy” scenario, as in Experiment 1. To what can this task-dependence be attributed? One likelihood is related to differences in the use of visual feedback control across the two scenarios. Several off-axis head rotations must be performed in the Trench scenario. Fine control of the vehicle when traversing the trench required large amounts of yaw head movements while the head was pitched forwards. It is at these moments when sickness-related sensory errors are most likely, due to so-called pseudo-coriolis effects (Bles 1988). As such, dFOV limits the nauseating stimuli at key moments, while allowing a greater field-of-view at other times.

This finding mirrors previous research showing that time-coupling vestibular stimulation to visual accelerations alleviates sickness more efficiently than when vestibular stimuli are delivered at random times (Weech et al. 2018a). These results can be considered consistent with computational accounts of multisensory integration, where the process by which time-coincident cues are combined to impact the sensory decision-making process has been well modeled in Bayesian (Knill & Pouget 2004) and maximum-likelihood terms (Parise et al. 2012).

## 5. General Discussion

In two user studies, we investigated the effects of cybersickness mitigations on a MEWP motion base simulator. In Experiment 1 we implemented a harness that provided minimal stability support, but that allowed participants to gather subtle cues about their spatial orientation from body haptics. In Experiment 2 we removed the harness and instead used a dynamic field-of-view (dFOV) limiter that blacked-out the eccentric visual flow when participants were rotating the virtual vehicle at a rate that was greater than some small threshold of angular velocity.

Both studies provided evidence that the mitigations we used were effective. Effectiveness of each mitigation differed at an absolute level (see below, “Mitigation Strength”) but both were considerably more useful in a “bumpy” task (see below, “Limitations”).

Experiment 1 was designed to test if a loosely tethered harness offers valuable cues to help users re-orient themselves in physical space, thus protecting against cybersickness, and we found results that were partially consistent with this idea. The harness may have provided a stream of feedback-control relevant cues and—crucially— relieved the emesis-related brain areas that are in part responsible for generating and updating state estimates of the head and body. Even though disorientation scores were unaffected by the harness, nausea was markedly improved. If there were a stabilizing effect of harness-wearing, it did not penetrate perception, but rather it is likely that the effects were told at the level of the brainstem or vestibular cerebellum (i.e., unconscious processing areas). Given that this is one region where subtle body orientation cues are thought to be integrated (Green & Angelaki 2010), our data may be considered unsurprising.

Experiment 2 confirmed that limiting eccentric visual flow during rotations can alleviate sickness. Although the dFOV limiter was only minimally tuned before this study was executed, many participants were brought from ‘sick’ to ‘well’ by the mitigation, according to an SSQ-T threshold of 20 (Stanney et al. 1997). The increased comfort that we documented when using dFOV agrees with cybersickness models that state the sickness center in the brain accrues errors over time primarily from optic flow in the wide periphery. As such, limiting visual flow in these areas slows the build-up of sickness (Lee et al. 2017).

Motion sickness and cybersickness mitigations that target bodily stability are commonplace (Keshavarz 2016; Stoffregen et al. 2010). However, the harness that we applied here was expressly intended to provide a ‘cue’ that the user could integrate with other sensory information to derive a more reliable estimate of the state of their body in 3D space as they executed their task in VR. We considered that uncertainty in the self-motion and self-orientation resolution process was a key contributor to cybersickness, as discussed elsewhere (Weech et al. 2020a), and reducing this uncertainty was our central focus. Although our results support the hypothesis that the harness effectively provided this information, it could equally be claimed that the use of a harness simply provided added postural stability, and that stability itself is the key contributor to user comfort (Stoffregen & Riccio 1991; Stoffregen & Smart 1998). While we did not intend to ‘restrain’ the users with the harness, we cannot entirely rule out alternative explanations for our results, which is a common problem in the domain of cybersickness (Riccio & Stoffregen 1991; Weech et al. 2018b).

The dFOV manipulation we used here has been applied in experimental research elsewhere (Fernandes & Feiner 2016; Lim et al. 2021; Teixeira & Palmisano 2021), with several commercial products now adding such a visual modifier to their software (e.g., Google Earth VR). Our results reiterate the power of a dFOV limiter when navigating virtual spaces, especially when associated with imposed angular motion. Visually induced self-motion (vection) has long been linked to motion sickness and cybersickness (see Keshavarz et al. 2015). Peripheral flow cues are the principal stimulus to the self-motion system (Gibson 1961; Warren & Hannon 1988), and visual motion at the far eccentricities have long been known to generate strong vection sensations, with lower FOV stimuli provoking weaker vection (Brandt et al. 1973; Johansson 1977; Murovec et al. 2021). Murovec and colleagues (2021), for instance, report a robust improvement in subjective vection as display FOV increases for a rotating visual stimulus. Therefore, it is conceivable that our results might reflect that vection was reduced by dFOV (note, we did not obtain vection estimates in our experiments). It should be noted, however, that no effect of dFOV was seen on vection in other research (Teixeira & Palmisano 2021).

As well as agreeing with the initial hypothesis that optic flow is a strong contributor to cybersickness, the results could also be taken as support for the recent theory that cybersickness results from differences in the virtual and physical head states (DVP, Palmisano et al. 2021). Assuming that visual flow from wide eccentricities is the most valuable source of information about head position, the sensed discrepancy between physical and virtual head locations should be lowest when peripheral information is occluded. However, there appears to be some ambiguity here regarding this perspective’s predictions: If the *precision* of the estimate of the head position in space were to become lower via the application of a dFOV limiter, this might result in greater sickness rather than the lower level of sickness that we reported here. At present, the DVP theory requires further elaboration to speak to these results.

In future work it may be desirable to modify the extent of the dFOV manipulation to account for individual differences in the useful field of view (UFOV, Ball et al. 1988), the area from which individuals reliably extract task-relevant information. This could enable the use of a ‘stronger’ (i.e., lower transparency) dFOV modification in those that do not utilize peripheral visual cues as much as others. Individual differences in field dependence that have recently been implicated in cybersickness (Fantin et al. 2022) could also provide valuable targets when considering an individually tailored dFOV modifier.

### 5.1. Mitigation Strength

We found that the ‘stronger’ mitigation at reducing sickness was the dFOV limiter. Total sickness scores were reduced by up to 45%, compared to a gross reduction of up to 11% for the harness (more so for nausea than other dimensions of cybersickness). Note that we did not test the combination of the two mitigations here. It would be informative to assess whether there is additive or even multiplicative value of using both simultaneously.

### 5.2. Limitations

One limitation of this work is that neither mitigation substantially affected sickness in the Choose Incline scenario. From these data, it appears that each mitigation might focally improve certain cases of sickness, but that a multi-pronged approach is likely required for battling against the wide variety of sickening scenarios to which users can be exposed.

In both experiments, the analysis would have benefited from greater statistical power. The easiest way to achieve this would have been to increase our sample size, which was limited due to practical constraints. Another way to do this would be to carry out a covariate analysis, factoring-out certain factors that vary from person to person. The inclusion of intrinsic factors known to be associated with cybersickness such as VR/gaming experience, age, sex, and motion sickness history (Stanney et al. 2020; Weech et al. 2020b) as covariates in future analyses will generate partial effects that more clearly depict the specific effects of mitigations. This would be best performed with a large, diverse sample of participants.

Due to imposed limits related to the emerging COVID-19 pandemic, time constraints meant that there was no scope for running pilot studies on the optimal way to implement each mitigation. For instance, we used only one configuration of harness tie-off points (Fig 2) and no adjustment was possible for the height or physique of the operator. For the dFOV, only one motion threshold for activation was used, and only one contrast gradient could be tested in our study. Increased iteration on both mitigations in future work would be highly valuable.

Here, we did not assess whether task performance, presence, or vection were inhibited by the visual field modifier. It has been previously suggested that limiting FOV might corrupt skilled performance training (Polys et al. 2005). However, emerging evidence suggests very limited effects of such techniques on skills acquisition, as demonstrated in spatial learning paradigms (Adhanom et al. 2021). Other work has found no deleterious effect of dFOV on the feeling of presence in a VR task (Fernandes & Feiner 2016). Although there is one report of intact levels of vection when a dFOV limiter is used (Teixeira & Palmisano 2021), there is a positive relationship between FOV and vection strength (Brandt et al. 1973; Johansson 1977; Murovec et al. 2021). Gathering these additional data in future work would help to evaluate if dFOV interferes with information acquisition in operational environments, such as personnel training.

## 6. Conclusion

Cybersickness mitigations are widely used in the VR domain. Here, we revealed evidence about the performance level of mitigations that aimed to reduce user discomfort in field relevant operational training tasks on a motion base MEWP simulator. Both a lightly tethered harness, and a dynamic field-of-view limiter, reduced sickness to varying extents—with the greatest impact observed for the field-of-view modification. Although the mitigations here were put to bear in small sample user studies, we gathered encouraging evidence that can be added to in future optimizations of these techniques.

## 7. Acknowledgements

This research was supported by funding from Mitacs through the Accelerate Fellowship program.

